# Pre-stimulus feedback connectivity biases the content of visual experiences

**DOI:** 10.1101/437152

**Authors:** Elie Rassi, Andreas Wutz, Nadia Müller-Voggel, Nathan Weisz

## Abstract

Ongoing fluctuations in neural excitability and in network-wide activity patterns before stimulus onset have been proposed to underlie variability in near-threshold stimulus detection paradigms, i.e. whether an object is perceived or not. Here, we investigated the impact of pre-stimulus neural fluctuations on the content of perception, i.e. whether one or another object is perceived. We recorded neural activity with magnetoencephalography before and while participants briefly viewed an ambiguous image, the Rubin face/vase illusion, and required them to report their perceived interpretation on each trial. Using multivariate pattern analysis, we showed robust decoding of the perceptual report during the post-stimulus period. Applying source localization to the classifier weights suggested early recruitment of V1 and ~160 ms recruitment of category-sensitive FFA. These post-stimulus effects were accompanied by stronger oscillatory power in the gamma frequency band for face vs vase reports. In pre-stimulus intervals, we found no differences in oscillatory power between face vs. vase reports neither in V1 nor in FFA, indicating similar levels of neural excitability. Despite this, we found stronger connectivity between V1 and FFA prior to face reports for low-frequency oscillations. Specifically, the strength of pre-stimulus feedback connectivity (i.e. Granger causality) from FFA to V1 predicted not only the category of the upcoming percept, but also the strength of post-stimulus neural activity associated with the percept. Our work shows that pre-stimulus network states can help shape future processing in category-sensitive brain regions and in this way bias the content of visual experiences.

## Introduction

Ongoing fluctuations in neural activity interact with perceptual and cognitive processes. They help explain why repetitions of the same physical stimuli elicit different percepts and responses from trial to trial in both animals (1) and humans (2–5). Both local excitability changes in task-relevant sensory regions (6, 7) and neural network connectivity patterns have been shown to underlie trial-by-trial fluctuations in perception (8–11).

The paradigms to study the impact of ongoing neural activity on perception typically involve near-threshold detection and discrimination tasks, in which pre-stimulus neural fluctuations influence the perceptual fate of stimuli, for example whether an object is seen (“Hit”) or not (“Miss”; e.g. (8, 10–19)). Beyond mere stimulus detection and discrimination, one of the visual system’s essential functions is to identify and categorize objects and, in this way, construct the content of visual experiences (19–21). Indeed, neural correlates of object perception and categorization have been shown to rely on the information flow between occipital and inferior temporal cortical regions (22–24). Here, we focus on the impact of neural excitability and connectivity patterns before stimulus onset on the content of perceptual operations.

Bi-stable perception paradigms are uniquely suited to address this question (25). In such paradigms, the brain is conflicted between multiple possible interpretations of visual content. Typical examples include the Rubin’s face/vase stimulus (26), the Necker cube (27), and binocular rivalry ((28); as cited in (29)). Recent evidence from fMRI studies has shown that rivalry between two competing percepts is resolved relatively early in the visual hierarchy (e.g. (30, 31)), such as in category-sensitive inferior temporal lobe regions ((32); but see (33) and (34) for fMRI and electrophysiological evidence showing an influence of parietal and frontal cortices; and see (35) for a recent review). In particular for the Rubin’s face/vase illusion, greater BOLD activity has been observed in the fusiform face area (FFA) when participants reported seeing a face rather than a vase (36). Importantly, this BOLD increase in FFA was also observed prior to stimulus onset (37), possibly because the pre-stimulus brain state biased perception towards the “face interpretation”. Still, a more comprehensive, mechanistic account requires means to simultaneously measure neural activity in multiple cortical areas with high temporal resolution, in order to map out the cortical hubs and their inter-areal information flow before and during perception of an ambiguous stimulus. For example, enhanced BOLD activity in FFA could be a consequence of either increased feedforward activity from earlier visual regions, or of increased feedback activity to earlier visual regions.

In the current study, we used a similar Rubin face/vase paradigm as the aforementioned fMRI study (37). Advancing on previous work, we thoroughly characterized neural activity and connectivity patterns with high temporal resolution prior to and during perception of the ambiguous Rubin stimulus by means of magnetoencephalography (MEG). We hypothesized that category-sensitive processing region (here FFA) should exhibit differential pre-stimulus connectivity patterns preceding subsequent face vs. vase reports. Based on the “Windows to Consciousness” framework (11, 38), fluctuating connectivity levels of sensory regions shape upcoming stimulus processing (i.e. whether a stimulus is perceived or not). We extended these predictions to visual object perception and investigated whether categorical responses to the content of the Rubin stimulus were biased by local excitability, feedforward connectivity, or feedback connectivity between primary visual cortex and FFA.

## Results

Twenty participants took part in the MEG experiment. We showed them the Rubin’s face / vase stimulus briefly and asked them to report whether they had seen faces or a vase on each trial (see SI Methods for details). Vase and face reports were equally likely (Face mean: 49.9 %; SD: 12.47%; range 22.6% to 84.8%; t-test against chance (50%) *t(19)* = 0.04, *p* = .97). To ascertain that the reported perception was stochastic trial-by-trial, we analysed the sequences of reported percepts by binning the trials into a range of 0 to 10 repetitions. A binomial distribution accounted well for the binned data for both vase and face trials (goodness-of-fit R^2^ = 0.96 for face, R^2^ = 0.98 for vase) indicative of no systematic reporting of either percept. That is, during each trial a participant was equally likely to report a face or vase irrespective of the previous trial.

In a first step, we aimed to extract category-specific information from the recorded MEG data to see whether source localization of this information would yield the ROIs found in previous work (i.e. V1 and FFA), and to later use this information to link pre- and post-stimulus neural activity. For this purpose, we trained a classifier in a cross-validation approach and decoded face vs vase reports from the MEG sensor-level data (magnetometers and gradiometers). The analysis was shifted over time on a sample-by-sample basis, yielding temporally resolved decoding results shown in **Fig. 1A**. Decoding performance (operationalized as Area Under Curve, AUC) gradually increased following stimulus onset and reached a peak close to the offset of the stimulus mask, the event which prompted the response query. From there on, decoding performance gradually decreased, reaching chance level after approximately 700 ms. Decoding accuracy was significantly above chance after 100 ms (*p*_cluster_ = 9.9990e-05; tested over the first 350 ms after stimulus onset to exclude the response epoch).

**Figure 1:**
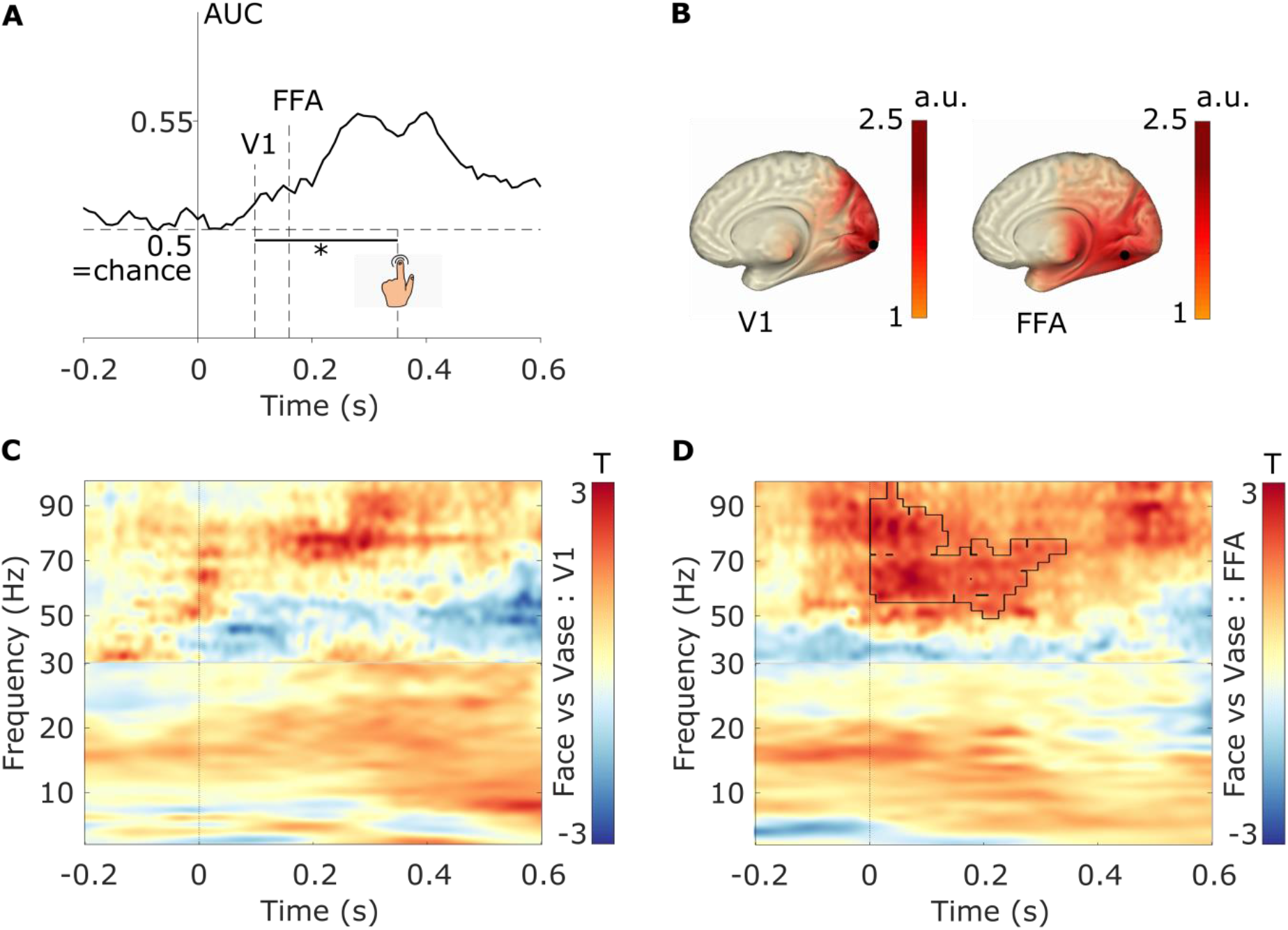
Post-stimulus MEG data contains category-sensitive information with respect to the processing of the Rubin vase stimulus. A) Temporal decoding of face vs vase reports. * represents p = .0001 significance of decoding accuracy (t-test vs chance) starting 100 ms post-stimulus. B) Unmasked activation maps resulting from the source reconstruction of the classifier weights (in arbitrary units a.u.), applying a procedure proposed by Marti and Dehaene (2017), at different time points, suggesting temporally changing informative regions (V1 around 100 ms and FFA around 160 ms after stimulus onset). C) Time-frequency contrast in V1 (face vs vase reports). Colors represent smoothed T-values obtained from cluster-based permutation testing of the contrast (face – vase; ns). D) Time-frequency contrast in FFA (face vs vase reports). Colors represent smoothed T-values obtained from cluster-based permutation testing of the contrast (face – vase; *p*_cluster_ = .029). Black lines surround the time-frequency gamma-range cluster that drove the significant statistical difference.

To find the brain regions that provided informative activity, we adapted a previously reported approach (39), which projects the classifier weights from sensor to source space (for sources see **Fig. 1B**). This analysis suggested that the brain regions that provide informative activity changed over time (**Fig. 1A**). At earlier (< 120 ms) time intervals, informative activity was predominantly localized in and around right V1 (centered on MNI coordinates: [12 −88 0] mm). In the subsequent time interval (120 – 200 ms), informative activity was predominantly localized in and around right FFA (centered on MNI coordinates: [28 −64 −4] mm). Although informative activity also spread to left V1 and FFA, the locations of maximum activity, which we used for subsequent analyses, were located in the right hemisphere. The location, lateralization, and timing of informative neural activity correspond well with reports on the spatiotemporal dynamics of face perception (40, 41). For all subsequent analysis, we used the source-reconstructed data from V1 and FFA.

Next, we performed time-frequency analysis in FFA after stimulus onset to reveal the oscillatory patterns that contributed to the decoding results. We contrasted trials on which participants reported seeing a face vs. a vase and corrected for multiple time-frequency samples with a cluster-based permutation approach (42). We found that face reports showed enhanced post-stimulus gamma activity (*p*_cluster_ = .029; **Fig. 1D**) compared to vase reports, consistent with the functional role of gamma activity for visual perception and specifically for face perception (43, 44). Over time, this cluster covered the entire relevant post-stimulus time-range and peaked at around 40ms. In terms of frequencies, the cluster covered a range between 48-93Hz and peaked between 60-70Hz. In the lower frequencies, there were no clusters in the time-frequency maps which contributed to the statistical effect (see **Fig. 1D**). We repeated the same analysis and contrast in V1 and found no statistical differences (no time-frequency clusters; **Fig. 1C**). Finally, we ran a sensor-wise time-frequency analysis, repeated the same contrast, and found no statistical differences on the whole-brain level (no time-frequency-sensor clusters). Overall, this analysis showed that perceiving the stimulus as face was accompanied by enhanced post-stimulus gamma activity in FFA.

The MVPA analysis yielded favorable ROIs to test whether pre-stimulus connectivity dynamics between early visual regions (V1) and later category-sensitive regions (FFA) bias the report of upcoming subjective percepts (see **Fig. 1A**). First, we focused on oscillatory power as an index of local excitability in these regions and tested whether excitability alone predicted the reported categories of upcoming stimuli. Oscillations reflect rhythmic changes in the activity of neural populations and thus reflect phases of high and low excitability (45). Cluster-based permutation testing revealed no statistical differences in pre-stimulus oscillatory power between face and vase trials, neither in V1 nor in FFA (**Fig. 2A**; shaded error regions represent the standard error of the mean for within-subjects designs (46)). Nevertheless, the power spectra in both conditions showed that pre-stimulus oscillatory activity was largely restricted to lower frequencies (5 to 25 Hz, see **Fig. 2A**) with a clear peak in the alpha range (~10 Hz). Since frequency-domain measures of connectivity (such as coherence or Granger causality) assume underlying oscillatory activity (i.e. oscillations with high power), we restricted statistical testing for the subsequent connectivity analyses to this frequency range.

**Figure 2:**
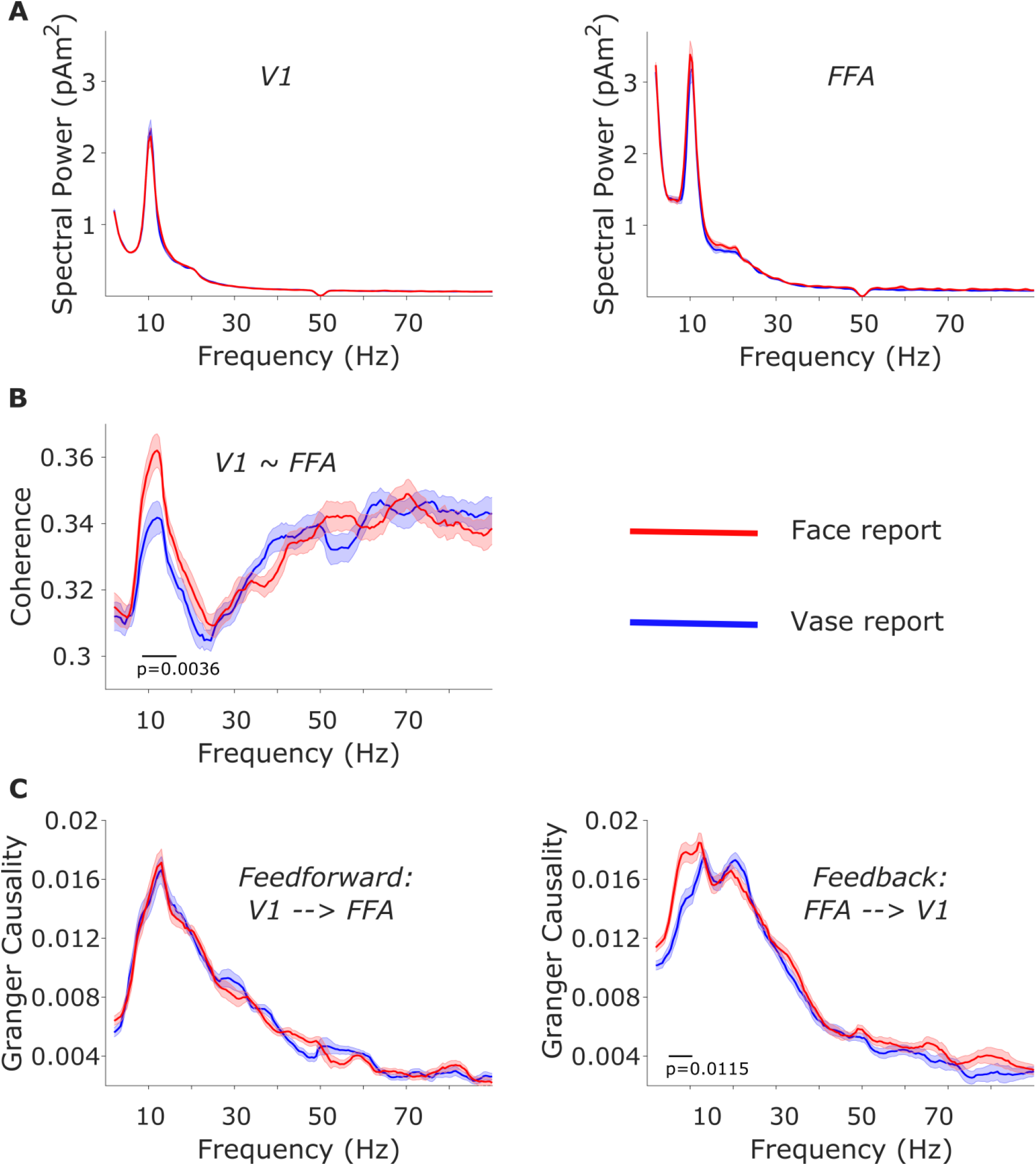
Pre-stimulus MEG connectivity is predictive of upcoming perceptual decision. Shaded error regions represent the standard error of the mean for within-subject designs (Morey, 2008). A) No statistical differences in pre-stimulus spectral power between face and vase trials in either V1 (left) or FFA (right). B) Compared to vase trials, face trials show increased pre-stimulus coherence between V1 and FFA in the alpha and beta frequency ranges. C) Compared to vase trials, face trials show increased pre-stimulus feedback connectivity from FFA to V1 in the alpha range (right), but no differences in pre-stimulus feedforward connectivity from V1 to FFA (left).

Next, we focused on pre-stimulus connectivity between V1 and FFA. Specifically, we hypothesized that increased pre-stimulus coherence between V1 and FFA would precede face reports. A cluster-based permutation test in the frequency range of 5 to 25 Hz revealed that pre-stimulus coherence between V1 and FFA was significantly greater on face vs. vase trials (p_cluster_ = .0036). This increase was most pronounced in a cluster of frequencies ranging from 8.5 to 16.5 Hz (**Fig. 2B**). To control for spurious coherence as a result of field spread (47), which might explain the high-frequency noise in **Fig. 2B**, we repeated the aforementioned analysis using the imaginary part of coherency (48). We obtained qualitatively and quantitatively similar results but with far less high-frequency.

To further characterize the observed connectivity effect, we used Granger causality to resolve the question of whether increased connectivity prior to face reports represented an increased feedforward drive from V1 to FFA, or an increased feedback drive from FFA to V1. We contrasted face and vase trials separately for the feedforward and feedback directions. The cluster-based permutation test revealed no statistical differences between face and vase reports in the pre-stimulus Granger causality estimates in the feedforward direction (V1 to FFA; **Fig. 2C, *left***); however, for feedback-connectivity we found significantly greater prestimulus Granger causality estimates during face trials compared to vase trials (FFA to V1, p_cluster_ = .0115). This increase was most pronounced in a cluster of frequencies ranging from 5 to 10.5 Hz (**Fig. 2C, *right***). The directionalities of the Granger estimates were reversed for time-reversed data (that is, the feedforward Granger estimates of the original data resembled the feedback Granger estimates of the time-reversed data, and vice versa), thereby confirming our results (49). Given the inter-individual variability in participants’ behavioral reports (22.6% to 84.8% face reports), we were concerned that the Granger results might reflect some participants’ predispositions to report one or the other percept. However, we found no correlation between the Granger strength and the report percentages (r=.22, p=.35), making this possibility unlikely. In sum, we show that face reports (vs. vase reports) were preceded by increased connectivity between V1 and FFA, and that this relative connectivity increase was predominantly driven by an increase in feedback connectivity (FFA to V1).

Finally, we focused on the relationship between pre-stimulus connectivity and post-stimulus activity. We extracted for each participant the maximum decoding accuracy (AUC), FFA gamma-band effect, and pre-stimulus feedback connectivity. The maximum FFA gamma effect (max. face – vase power over time and frequencies) and maximum decoding accuracy were correlated (r=.58, p = .008; **Fig. 3C**), despite the gamma band having been excluded from the frequency range that went into the decoder. Importantly, we found that maximum pre-stimulus feedback connectivity was correlated with both the maximum gamma effect (r = .57, p = .009; **Fig. 3A**) and maximum decoding accuracy (r = .48, p = .034, **Fig. 3B**). In sum, we found that pre-stimulus feedback connectivity strength predicted not only the category of the upcoming percept, but also the strength of post-stimulus neural activity associated with the percept.

**Figure 3:**
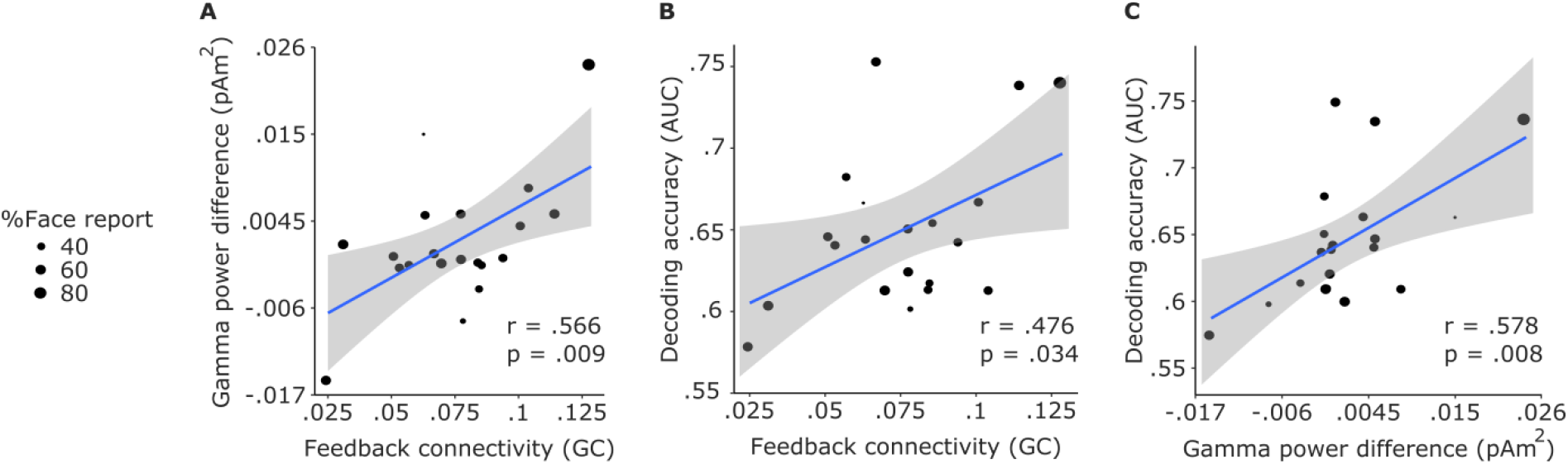
Pre-stimulus connectivity is correlated with post-stimulus activity across participants. r values represent Pearson’s correlation coefficients. Shaded areas represent 95% confidence intervals. A) Maximum pre-stimulus feedback Granger causality estimates are correlated with maximum post-stimulus gamma differences (face – vase). B) Maximum pre-stimulus feedback Granger causality estimates are correlated with maximum post-stimulus decoding (AUC) scores. C) Maximum post-stimulus gamma differences (face – vase) are correlated with maximum post-stimulus decoding (AUC) scores.

## Discussion

While most studies that investigated the effects of pre-stimulus activity on perception were concerned with determining the requisites of successfully detecting stimuli at perceptual threshold (near-threshold paradigms; e.g. (11)), our main interest was with the requisites of perceiving one or another content of perception. We found that prior to the Rubin face/vase stimulus onset, FFA was more strongly connected to V1 when face rather than vase was subsequently reported, specifically in the feedback direction of FFA to V1. Connectivity between these two regions was concentrated in the alpha and beta frequency bands (around 5 to 25 Hz). Further, pre-stimulus feedback connectivity strength was correlated with post-stimulus neural activity strength as well as decoding accuracy. Taken together, our findings suggest that fluctuations in neural activity in the absence of stimulation can bias the perceptual content of subsequently presented stimuli.

The connectivity pathway we’ve identified, specifically the involvement of FFA, is well in line with works that have localized face responses using ambiguous stimuli (eg: (36, 50)). While this particular pathway is likely specific to face stimuli, the involvement of functionally specialized extrastriate regions in the subjective perception of ambiguous stimuli is firmly established (51). Indeed, processing semantic content typically relates to ventral stream activity, so this activity is also expected to play a crucial role in perceiving ambiguous images of semantic content, such as the Rubin vase image (52). That the connectivity pathway is in the feedback direction and in the lower frequencies is also well in line with the finding that alpha/beta oscillations subserve feedback connectivity among human (53) and monkey (54) visual cortical areas. Additionally, occipital alpha oscillations have been shown to predict the persistence of bistable perception (55), and percept-dependent changes in occipital oscillatory activity have been suggested to reflect top-down modulations of V1 by extrastriate areas (56). Our findings therefore suggest that in the absence of visual stimulation, mechanisms that mimic known dynamics of unambiguous as well as ambiguous visual object perception are at play.

MEG studies on face perception have reported gamma responses to faces starting 100 ms after stimulus onset (e.g. (57, 58)), yet we observed a peak gamma response at 40 ms. But while these studies typically compare responses to faces with non-face stimuli and apply baseline corrections to their data, we compared responses to the same exact stimulus and did not apply any baseline corrections such that we could observe potential pre- and peri-stimulus effects. The early latency of our gamma effect is in line with recent MEG work showing differential 40ms evoked responses to forms (59). The 40 ms response was also reported for face stimuli and localized in FFA and other visual areas (60), indicating parallelism in visual input streams and challenging the commonly held supposition that V1 is the sole entry point for visual input (61, 62). That our gamma cluster starts at stimulus onset, and is to some extent observable even before, suggests that it likely reflects an interaction between pre-stimulus anticipatory processes (in which gamma oscillations have also been implicated; eg: (14, 63, 64)) and very early sensory processes. Modelling work has suggested that gamma oscillations enhance neural network responsiveness (65), a mechanism possibly mediating our pre- and post-stimulus effects. That this gamma effect was correlated with both the strength of pre-stimulus feedback (**Fig. 3A**) and post-stimulus decoding accuracy (**Fig. 3C**) lends support to this interpretation. Given that the pre-stimulus connectivity pathway was in the feedback direction from FFA to V1, one might additionally hypothesize that the strength of feedback connectivity correlates with V1 gamma activity. But V1 gamma modulations have been shown to depend on stimulus features (e.g. (66)), and the stimulus was unchanged throughout our experiment. Indeed, we did not observe any gamma effects in V1 (**Fig. 1C**), so we could not test this hypothesis.

A recent fMRI study employing the Rubin face-vase stimulus (37) found that pre- and post-stimulus neural activity was pronounced in the FFA and interpreted the observed pre-stimulus BOLD signal differences as differences in baseline excitability. Indeed, this interpretation is consistent with a large body of work that shows that alpha band activity in task-sensitive sensory regions, an index of neuronal excitability in those regions, predicts behavioral outcome (8, 10, 11, 67). Yet, we found no differences in pre-stimulus alpha activity that could account for the behavioral outcome. Local excitability as indexed by alpha oscillations might therefore be behaviorally relevant in near-threshold cases, but not in cases where the stimuli are supra-threshold and the task requires object perception rather than stimulus detection or discrimination. Taken together, these results show that measures of local activity paint an incomplete picture of the underlying dynamics of object perception, and that the connectivity between regions of interest must be considered for a more comprehensive account.

Given the nature of this and similar experiments (e.g. (37)), it is difficult to pinpoint distinct cognitive processes to our effects with certainty. However, our findings are well in line with predictive processing notions that hierarchically downstream regions predict activity in upstream areas via feedback connections. The reported frequency band conforms to the assumptions and findings of this framework (68, 69). Since this connectivity effect precedes the presentation of the ambiguous stimulus, an interpretation along the lines of anticipatory predictive processes appears promising (70). Indeed, our findings add to a recent and fast-growing literature converging towards the idea that low-frequency oscillations carry top-down context (71), category information (72), anticipation (73), and expectations or predictions (74, 75). That cognitive, top-down influences come to play leaves open the possibility that the reported connectivity effects are not strictly spontaneous, and might be voluntarily driven to some extent. This possibility cannot be entirely ruled out, although it is unlikely given our design (short, temporally difficult-to-predict inter-stimulus intervals of 1-1.8 s) and behavioural analysis which ruled out systematic reports of one percept. Relatedly, an alternative explanation could be that stronger connectivity relates to a stronger predisposition to perceive a face. However, connectivity strength was not correlated with the percentage of face reports, making this interpretation unlikely too.

Our results are in line with the Windows to Consciousness framework which emphasizes the influence of pre-established connectivity patterns of relevant sensory regions to downstream processing regions on upcoming perceptual processing (11, 38). We offer a mechanistic account defined in time, space, oscillatory frequency, and directional connectivity. Our account proposes a key role of pre-stimulus neural fluctuations in activating connectivity pathways and biasing categorical percepts. Specifically, pre-stimulus feedback connectivity in the alpha range from FFA to V1 represents such a connectivity pathway that biases towards face perception of the Rubin face/vase stimulus.

## Conclusion

By recording MEG signals at high temporal resolution before and while people were exposed to an ambiguous stimulus, the Rubin face/vase illusion, we showed that the content of visual perception is critically shaped by the ongoing network states, in this case feedback alpha-band connectivity between face-sensitive FFA and early visual cortex. Our work bridges object perception-related pre- and post-stimulus effects and shows how a pre-stimulus network state can shape future processing in a category-sensitive brain region.

## Materials and Methods

20 volunteers participated in this MEG experiment. At the beginning of each trial, a fixation cross would appear at the centre of the screen for 1 to 1.8 s. After this jittered period, the Rubin vase picture would appear at the centre of the screen for 150 ms (**Fig. 4**). A mask stimulus would then appear for 200 ms, after which we asked participants to report whether they saw the face or the vase. The experiment consisted of 400 trials in total.

**Figure 4:**
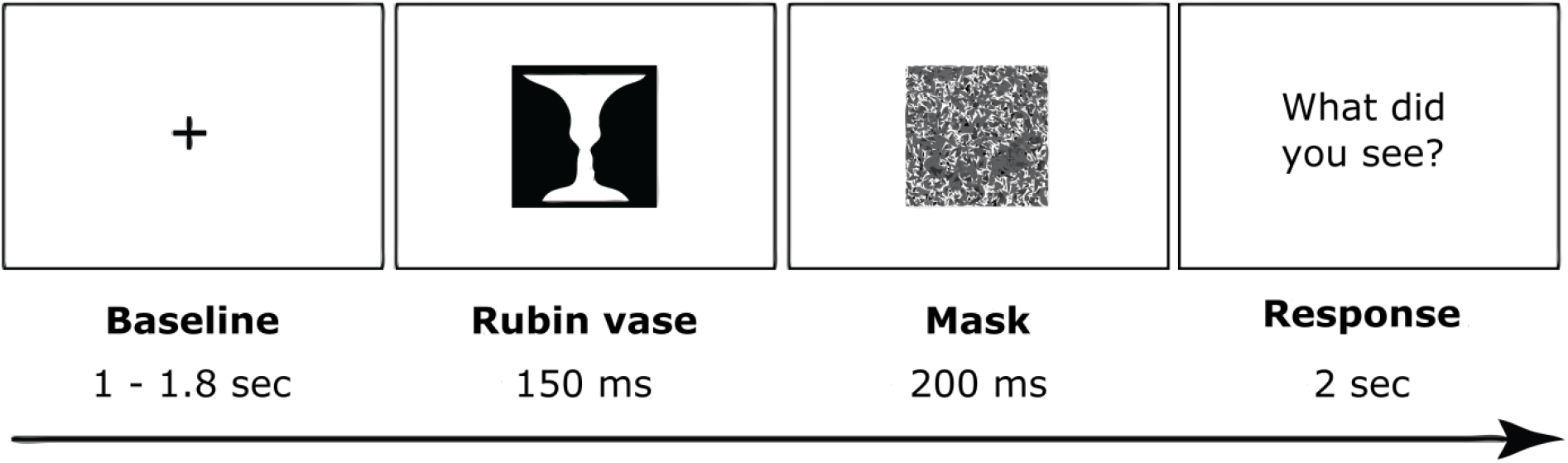
Trial structure.

To test for the stochastic nature of the response, we binned the data for each participant according to how many trials in a row they responded with the same perceptual report. We broke this down in 11 bins with 0 repetitions to 10 repetitions, averaged the number of repetitions within each bin across participants, and then fit the averaged data to a binomial distribution across the 11 bins before calculating goodness-of-fit.

We performed the decoding analysis on the broadband 1-33 Hz time-domain signal. We implemented a 4-fold cross-validation procedure within each subject. The analysis was shifted over time on a sample-by-sample basis. For each time-point at each sensor, we Z-normalized the MEG data, trained a Logistic Regression classifier on three folds, and tested on the left-out fold. To find out which brain regions contributed to above chance decoding performance the most, we used the classifier weights that the classifier used to separate face from vase reports and projected them into source space (39). Finally, we averaged the source-level weights across the intervals 50 to 120 ms and 120 to 200 ms and applied a 95%-max threshold to mask our ROIs.

We performed the post-stimulus time-frequency analysis on V1, FFA, and the whole-brain average in source space. We estimated power using multitaper Fast Fourier Transform (FFT) with discrete prolate spheroidal sequences (dpss) tapers (76). We calculated power, coherence, and Nonparametric Granger causality (77) in the pre-stimulus period between FFA and V1 in source space. We used multi-taper frequency transformation to get Fourier coefficients in the pre-stimulus period (−1 to 0 s), after which we extracted power and computed coherence and bivariate Granger causality. This gave us separate estimates of connection strengths from FFA to V1 (“feedback”) and vice versa (“feedforward”). We repeated the same Granger causality analysis on time-reversed data, expecting reversals in the directionalities of the estimates to rule out spurious connectivity results (49).

We tested decoding performance against chance level (50%) using one-sided dependent-samples T-tests. For all remaining statistical analyses, we used nonparametric cluster permutation tests (42). We used 2-sided T-tests for the post-stimulus time-frequency contrasts and pre-stimulus power contrasts, and 1-sided T-tests for the coherence and feedforward and feedback connectivity contrasts, as we had hypothesized greater values of these measures on face trials compared to vase trials. We restricted the statistical testing window of coherence and Granger to the frequency window 5-25 Hz.

## Acknowledgements

This work was supported by FWF Austrian Science Fund, Imaging the Mind: Connectivity and Higher Cognitive Function, W 1233-G17 (to E.R.), European Research Council Grant WIN2CON, ERC StG 283404 (to N.W.), and FWF-Lise Meitner fellowship, M02496 (to A.W.). We thank Nicholas Peatfield for valuable contributions to this work.

## References

1. A. Arieli, A. Sterkin, A. Grinvald, & A. Aertsen, Dynamics of ongoing activity: explanation of the large variability in evoked cortical responses. Science 273, 1868–1871 (1996).

2. E. Basar, A. Gonder, & P. Ungan, Important relation between EEG and brain evoked potentials. II. A systems analysis of electrical signals from the human brain. Biol Cybern 25, 41–48 (1976).

3. E. Basar, A. Gonder, & P. Ungan, Important relation between EEG and brain evoked potentials. I. Resonance phenomena in subdural structures of the cat brain. Biol Cybern 25, 27–40 (1976).

4. M. Boly, E. Balteau, C. Schnakers, C. Degueldre, G. Moonen, A. Luxen, C. Phillips, P. Peigneux, P. Maquet, & S. Laureys, Baseline brain activity fluctuations predict somatosensory perception in humans. Proc Natl Acad Sci U S A 104, 12187–12192 (2007).

5. E. Rahn & E. Basar, Prestimulus EEG-activity strongly influences the auditory evoked vertex response: a new method for selective averaging. Int J Neurosci 69, 207–220 (1993).

6. H. Super, C. van der Togt, H. Spekreijse, & V. A. Lamme, Internal state of monkey primary visual cortex (V1) predicts figure-ground perception. J Neurosci 23, 3407–3414 (2003).

7. S. Sadaghiani, G. Hesselmann, & A. Kleinschmidt, Distributed and antagonistic contributions of ongoing activity fluctuations to auditory stimulus detection. J Neurosci 29, 13410–13417 (2009).

8. J. N. Frey, P. Ruhnau, S. Leske, M. Siegel, C. Braun, & N. Weisz, The Tactile Window to Consciousness is Characterized by Frequency-Specific Integration and Segregation of the Primary Somatosensory Cortex. Sci Rep 6, 20805 (2016).

9. E. Leonardelli, C. Braun, N. Weisz, C. Lithari, V. Occelli, & M. Zampini, Prestimulus oscillatory alpha power and connectivity patterns predispose perceptual integration of an audio and a tactile stimulus. Hum Brain Mapp 36, 3486–3498 (2015).

10. S. Leske, P. Ruhnau, J. Frey, C. Lithari, N. Muller, T. Hartmann, & N. Weisz, Prestimulus Network Integration of Auditory Cortex Predisposes Near-Threshold Perception Independently of Local Excitability. Cereb Cortex 25, 4898–4907 (2015).

11. N. Weisz, A. Wuhle, G. Monittola, G. Demarchi, J. Frey, T. Popov, & C. Braun, Prestimulus oscillatory power and connectivity patterns predispose conscious somatosensory perception. Proc Natl Acad Sci U S A 111, E417–425 (2014).

12. C. S. Y. Benwell, C. F. Tagliabue, D. Veniero, R. Cecere, S. Savazzi, & G. Thut, Prestimulus EEG Power Predicts Conscious Awareness But Not Objective Visual Performance. eNeuro 4, (2017).

13. T. Ergenoglu, T. Demiralp, Z. Bayraktaroglu, M. Ergen, H. Beydagi, & Y. Uresin, Alpha rhythm of the EEG modulates visual detection performance in humans. Brain Res Cogn Brain Res 20, 376–383 (2004).

14. S. Hanslmayr, A. Aslan, T. Staudigl, W. Klimesch, C. S. Herrmann, & K. H. Bauml, Prestimulus oscillations predict visual perception performance between and within subjects. Neuroimage 37, 1465–1473 (2007).

15. L. Iemi, M. Chaumon, S. M. Crouzet, & N. A. Busch, Spontaneous Neural Oscillations Bias Perception by Modulating Baseline Excitability. J Neurosci 37, 807–819 (2017).

16. D. Kaiser, N. N. Oosterhof, & M. V. Peelen, The Neural Dynamics of Attentional Selection in Natural Scenes. J Neurosci 36, 10522–10528 (2016).

17. H. van Dijk, J. M. Schoffelen, R. Oostenveld, & O. Jensen, Prestimulus oscillatory activity in the alpha band predicts visual discrimination ability. J Neurosci 28, 1816–1823 (2008).

18. A. Wutz & D. Melcher, The temporal window of individuation limits visual capacity. Front Psychol 5, 952 (2014).

19. A. Wutz, D. Melcher, & J. Samaha, Frequency modulation of neural oscillations according to visual task demands. Proc Natl Acad Sci U S A 115, 1346–1351 (2018).

20. N. K. Logothetis & D. L. Sheinberg, Visual object recognition. Annu Rev Neurosci 19, 577–621 (1996).

21. E. K. Miller, A. Nieder, D. J. Freedman, & J. D. Wallis, Neural correlates of categories and concepts. Curr Opin Neurobiol 13, 198–203 (2003).

22. D. J. Freedman, M. Riesenhuber, T. Poggio, & E. K. Miller, A comparison of primate prefrontal and inferior temporal cortices during visual categorization. J Neurosci 23, 5235–5246 (2003).

23. G. Kreiman, C. Koch, & I. Fried, Category-specific visual responses of single neurons in the human medial temporal lobe. Nat Neurosci 3, 946–953 (2000).

24. N. Sigala & N. K. Logothetis, Visual categorization shapes feature selectivity in the primate temporal cortex. Nature 415, 318–320 (2002).

25. R. Blake & N. Logothetis, Visual competition. Nat Rev Neurosci 3, 13–21 (2002).

26. E. Rubin, Synsoplevede figurer. (1915).

27. L. A. Necker, LXI. Observations on some remarkable optical phænomena seen in Switzerland; and on an optical phænomenon which occurs on viewing a figure of a crystal or geometrical solid. The London, Edinburgh, and Dublin Philosophical Magazine and Journal of Science 1, 329–337 (1832).

28. J. Porta, 1593 De Refractione. Optices Parte. Libri Novem (Naples: Salviani),

29. N. J. Wade, Descriptions of visual phenomena from Aristotle to Wheatstone. Perception 25, 1137–1175 (1996).

30. D. A. Leopold & N. K. Logothetis, Activity changes in early visual cortex reflect monkeys’ percepts during binocular rivalry. Nature 379, 549–553 (1996).

31. J. Zou, S. He, & P. Zhang, Binocular rivalry from invisible patterns. Proc Natl Acad Sci U S A 113, 8408–8413 (2016).

32. F. Tong, K. Nakayama, J. T. Vaughan, & N. Kanwisher, Binocular rivalry and visual awareness in human extrastriate cortex. Neuron 21, 753–759 (1998).

33. P. Sterzer & A. Kleinschmidt, A neural basis for inference in perceptual ambiguity. Proc Natl Acad Sci U S A 104, 323–328 (2007).

34. M. Vernet, A. K. Brem, F. Farzan, & A. Pascual-Leone, Synchronous and opposite roles of the parietal and prefrontal cortices in bistable perception: a double-coil TMS-EEG study. Cortex 64, 78–88 (2015).

35. J. Brascamp, P. Sterzer, R. Blake, & T. Knapen, Multistable Perception and the Role of the Frontoparietal Cortex in Perceptual Inference. Annu Rev Psychol 69, 77–103 (2018).

36. U. Hasson, T. Hendler, D. Ben Bashat, & R. Malach, Vase or face? A neural correlate of shape-selective grouping processes in the human brain. J Cogn Neurosci 13, 744–753 (2001).

37. G. Hesselmann, C. A. Kell, E. Eger, & A. Kleinschmidt, Spontaneous local variations in ongoing neural activity bias perceptual decisions. Proc Natl Acad Sci U S A 105, 10984–10989 (2008).

38. P. Ruhnau, A. Hauswald, & N. Weisz, Investigating ongoing brain oscillations and their influence on conscious perception - network states and the window to consciousness. Front Psychol 5, 1230 (2014).

39. S. Marti & S. Dehaene, Discrete and continuous mechanisms of temporal selection in rapid visual streams. Nat Commun 8, 1955 (2017).

40. K. Botzel, S. Schulze, & S. R. Stodieck, Scalp topography and analysis of intracranial sources of face-evoked potentials. Exp Brain Res 104, 135–143 (1995).

41. N. Kanwisher, J. McDermott, & M. M. Chun, The fusiform face area: a module in human extrastriate cortex specialized for face perception. J Neurosci 17, 4302–4311 (1997).

42. E. Maris & R. Oostenveld, Nonparametric statistical testing of EEG- and MEG-data. J Neurosci Methods 164, 177–190 (2007).

43. A. D. Engell & G. McCarthy, The relationship of gamma oscillations and face-specific ERPs recorded subdurally from occipitotemporal cortex. Cereb Cortex 21, 1213–1221 (2011).

44. L. Fisch, E. Privman, M. Ramot, M. Harel, Y. Nir, S. Kipervasser, F. Andelman, M. Y. Neufeld, U. Kramer, I. Fried, & R. Malach, Neural “ignition”: enhanced activation linked to perceptual awareness in human ventral stream visual cortex. Neuron 64, 562–574 (2009).

45. W. Klimesch, P. Sauseng, & S. Hanslmayr, EEG alpha oscillations: the inhibition-timing hypothesis. Brain Res Rev 53, 63–88 (2007).

46. R. D. Morey, Confidence intervals from normalized data: A correction to Cousineau (2005). reason 4, 61–64 (2008).

47. A. M. Bastos & J. M. Schoffelen, A Tutorial Review of Functional Connectivity Analysis Methods and Their Interpretational Pitfalls. Front Syst Neurosci 9, 175 (2015).

48. G. Nolte, O. Bai, L. Wheaton, Z. Mari, S. Vorbach, & M. Hallett, Identifying true brain interaction from EEG data using the imaginary part of coherency. Clin Neurophysiol 115, 2292–2307 (2004).

49. S. Haufe, V. V. Nikulin, K. R. Muller, & G. Nolte, A critical assessment of connectivity measures for EEG data: a simulation study. Neuroimage 64, 120–133 (2013).

50. P. Sterzer & G. Rees, A neural basis for percept stabilization in binocular rivalry. J Cogn Neurosci 20, 389–399 (2008).

51. P. Sterzer, A. Kleinschmidt, & G. Rees, The neural bases of multistable perception. Trends Cogn Sci 13, 310–318 (2009).

52. G. A. Rodríguez-Martínez & H. Castillo-Parra, Bistable perception: neural bases and usefulness in psychological research. International Journal of Psychological Research 11, 63–76 (2018).

53. G. Michalareas, J. Vezoli, S. van Pelt, J. M. Schoffelen, H. Kennedy, & P. Fries, Alpha-Beta and Gamma Rhythms Subserve Feedback and Feedforward Influences among Human Visual Cortical Areas. Neuron 89, 384–397 (2016).

54. T. van Kerkoerle, M. W. Self, B. Dagnino, M. A. Gariel-Mathis, J. Poort, C. van der Togt, & P. R. Roelfsema, Alpha and gamma oscillations characterize feedback and feedforward processing in monkey visual cortex. Proc Natl Acad Sci U S A 111, 14332–14341 (2014).

55. G. Piantoni, N. Romeijn, G. Gomez-Herrero, Y. D. Van Der Werf, & E. J. W. Van Someren, Alpha Power Predicts Persistence of Bistable Perception. Sci Rep 7, 5208 (2017).

56. L. Parkkonen, J. Andersson, M. Hamalainen, & R. Hari, Early visual brain areas reflect the percept of an ambiguous scene. Proc Natl Acad Sci U S A 105, 20500–20504 (2008).

57. G. Perry & K. D. Singh, Localizing evoked and induced responses to faces using magnetoencephalography. Eur J Neurosci 39, 1517–1527 (2014).

58. S. Uono, W. Sato, T. Kochiyama, Y. Kubota, R. Sawada, S. Yoshimura, & M. Toichi, Time course of gamma-band oscillation associated with face processing in the inferior occipital gyrus and fusiform gyrus: A combined fMRI and MEG study. Hum Brain Mapp 38, 2067–2079 (2017).

59. Y. Shigihara & S. Zeki, Parallelism in the brain’s visual form system. Eur J Neurosci 38, 3712–3720 (2013).

60. Y. Shigihara & S. Zeki, Parallel processing of face and house stimuli by V1 and specialized visual areas: a magnetoencephalographic (MEG) study. Front Hum Neurosci 8, 901 (2014).

61. S. Zeki, A massively asynchronous, parallel brain. Philos Trans R Soc Lond B Biol Sci 370, (2015).

62. S. Zeki, Multiple asynchronous stimulus- and task-dependent hierarchies (STDH) within the visual brain’s parallel processing systems. Eur J Neurosci 44, 2515–2527 (2016).

63. A. K. Engel, P. Fries, & W. Singer, Dynamic predictions: oscillations and synchrony in top-down processing. Nat Rev Neurosci 2, 704–716 (2001).

64. P. Fries, J. H. Reynolds, A. E. Rorie, & R. Desimone, Modulation of oscillatory neuronal synchronization by selective visual attention. Science 291, 1560–1563 (2001).

65. S. B. Paik, T. Kumar, & D. A. Glaser, Spontaneous local gamma oscillation selectively enhances neural network responsiveness. PLoS Comput Biol 5, e1000342 (2009).

66. E. V. Orekhova, O. V. Sysoeva, J. F. Schneiderman, S. Lundstrom, I. A. Galuta, D. E. Goiaeva, A. O. Prokofyev, B. Riaz, C. Keeler, N. Hadjikhani, C. Gillberg, & T. A. Stroganova, Input-dependent modulation of MEG gamma oscillations reflects gain control in the visual cortex. Sci Rep 8, 8451 (2018).

67. V. Wyart & C. Tallon-Baudry, How ongoing fluctuations in human visual cortex predict perceptual awareness: baseline shift versus decision bias. J Neurosci 29, 8715–8725 (2009).

68. A. M. Bastos, W. M. Usrey, R. A. Adams, G. R. Mangun, P. Fries, & K. J. Friston, Canonical microcircuits for predictive coding. Neuron 76, 695–711 (2012).

69. A. M. Bastos, J. Vezoli, C. A. Bosman, J. M. Schoffelen, R. Oostenveld, J. R. Dowdall, P. De Weerd, H. Kennedy, & P. Fries, Visual areas exert feedforward and feedback influences through distinct frequency channels. Neuron 85, 390–401 (2015).

70. F. P. de Lange, M. Heilbron, & P. Kok, How Do Expectations Shape Perception? Trends Cogn Sci 22, 764–779 (2018).

71. R. F. Helfrich, M. Huang, G. Wilson, & R. T. Knight, Prefrontal cortex modulates posterior alpha oscillations during top-down guided visual perception. Proc Natl Acad Sci U S A 114, 9457–9462 (2017).

72. C. G. Richter, R. Coppola, & S. L. Bressler, Top-down beta oscillatory signaling conveys behavioral context in early visual cortex. Sci Rep 8, 6991 (2018).

73. R. Solis-Vivanco, O. Jensen, & M. Bonnefond, Top-Down Control of Alpha Phase Adjustment in Anticipation of Temporally Predictable Visual Stimuli. J Cogn Neurosci 30, 1157–1169 (2018).

74. A. Mayer, C. M. Schwiedrzik, M. Wibral, W. Singer, & L. Melloni, Expecting to See a Letter: Alpha Oscillations as Carriers of Top-Down Sensory Predictions. Cereb Cortex 26, 3146–3160 (2016).

75. J. Samaha, P. Bauer, S. Cimaroli, & B. R. Postle, Top-down control of the phase of alpha-band oscillations as a mechanism for temporal prediction. Proc Natl Acad Sci U S A 112, 8439–8444 (2015).

76. P. P. Mitra & B. Pesaran, Analysis of dynamic brain imaging data. Biophys J 76, 691–708 (1999).

77. M. Dhamala, G. Rangarajan, & M. Ding, Analyzing information flow in brain networks with nonparametric Granger causality. Neuroimage 41, 354–362 (2008).

